# A spiking neural network model of cortical intraregional metastability

**DOI:** 10.1101/2022.09.28.509893

**Authors:** Siva Venkadesh, Asmir Shaikh, Heman Shakeri, Ernest Barreto, John D. Van Horn

## Abstract

Transient synchronization of bursting activity in neural networks, which occurs in patterns of metastable phase relationships between neurons, is a notable feature of network dynamics observed *in vivo*. However, the mechanisms that contribute to this dynamical complexity in neural circuits are not well understood. Local circuits in cortical regions consist of populations of neurons with diverse intrinsic oscillatory features. In this study, we numerically show that the phenomenon of transient synchronization, also referred to as metastability, emerges in an inhibitory neural population when the neurons’ intrinsic fast-spiking dynamics are appropriately modulated by slower inputs from an excitatory neural population. Using a compact model of a mesoscopic-scale network consisting of excitatory pyramidal and inhibitory fast-spiking neurons, our work demonstrates a relationship between the frequency of neural oscillations and the features of emergent metastability. In addition, a novel metric is formulated to characterize collective transitions in metastable networks. Finally, we discuss a blueprint to model the whole-brain resting-state dynamics using our scalable representation of intraregional network metastability.

## 1. Introduction

Computational modeling of neuronal dynamics which incorporates mesoscopic-level connectivity features can potentially offer powerful frameworks for investigating neural network mechanisms [1,2]. However, a scalable network representation of key neurodynamical features at the mesoscopic level is currently lacking. This is primarily due to limited theoretical accounts for the emergent dynamical features in a network in terms of its constituent mechanisms. A simplified model describing the complexities of between- and within-group neuronal interactions is necessary to advance our multiscale understanding of neural systems in typical and atypical conditions [3].

An experimentally observable complexity of *in vivo* networks is the metastable attractor dynamics at the level of individual neurons. More specifically, a neuron can exhibit bursting oscillations that transiently synchronize with the oscillations of other neurons, and these synchronizations can occur at different phases [4–9]. In other words, biological neural networks can realize a range of attractor states that are locally observable through different phase-locking modes between individual neurons. Such transient attractor dynamics are also referred to as metastability or itinerancy [10], and this phenomenon is hypothesized to be a necessary physical property underlying the coordinated dynamics between spatially distant neural populations [11]. Moreover, a moment of conscious experience is hypothesized to exist irresolvably (i.e., indivisible in time) for the duration between successive attractor transitions in the brain [12].

The mathematical conditions for the emergence of transient dynamics have been previously discussed [13]. More recently, conduction delays between brain areas were associated with metastable oscillations on a macroscopic spatial scale [14]. However, the biophysical features of neural networks underlying metastability at the level of neurons have not been fully elucidated. While a previous work reported metastable attractor dynamics among intrinsically bursting neurons [10], it was unclear if metastability could also be realized in neural populations in which bursting dynamics are induced externally. Here, we explore the emergence and characteristics of metastability in a fast-spiking interneuron population that exhibits bursting dynamics due to driving inputs from a population of slower pyramidal neurons. We also examine the quantitative characteristics of collective transitions between attractor states realized in different metastable networks. Finally, we discuss potential applications of our work in modeling the whole-brain resting-state dynamics to understand network physiology in neurodegenerative disorders.

## 2. Methods

Networks consisting of 100 pyramidal neurons and 50 fast-spiking interneurons were constructed using Izhikevich neurons [15] (Equations 1 – 3). The parameters *a, b, c*, and *d* (Equations 2 – 3)

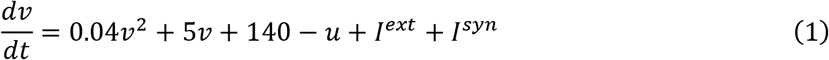

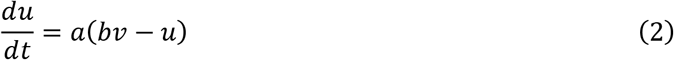

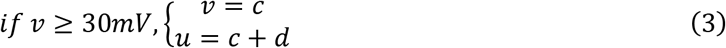

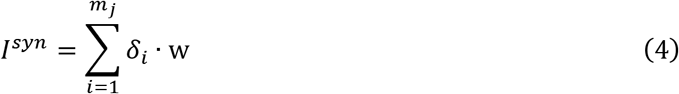

were selected to match the intrinsic dynamics of each type of neuron. Specifically, these were set to 0.02, 0.2, −65, and 8 respectively for the pyramidal neurons, and 0.1, 0.2, −65 and 2 for the fast-spiking interneurons [15].

Network connections were configured as illustrated in **Fig. 1A**. Inter- and intra-population connections were specified using pairwise connection probabilities between individual neurons. We studied the network behavior for a complete range of connection probabilities. Post-synaptic effects were specified using an instantaneous pulse-coupling scheme (Equation – 4), where each presynaptic spike either increased (excitatory) or decreased (inhibitory) the postsynaptic current by the weight parameter w (+0.3 for excitatory connections and −0.3 for inhibitory connections). In Equation – 4, *δ_i_* is a function which equals one when neuron *i* spikes and is zero otherwise. The number of neurons and the weight parameters were chosen to allow the pyramidal population to induce sustained bursting activity in the fast-spiking interneuron population for a range of external input currents in the former (*I^ext^* in Equation – 1). *I^ext^* was always set to zero for the fast-spiking interneurons. All networks were simulated using Brian2 [16] for a duration of 10s with a timestep of 0.1ms. Following the network simulation, a low-pass filter was applied to the voltage signal of each neuron (*v* in Equation – 1) to extract the bursting oscillations without the individual spikes (**Fig. 1C top**). Then, instantaneous phases of the bursting oscillations were assigned to each neuron via Hilbert transform. **Fig. 1C bottom** shows two representative examples.

**Figure 1.**
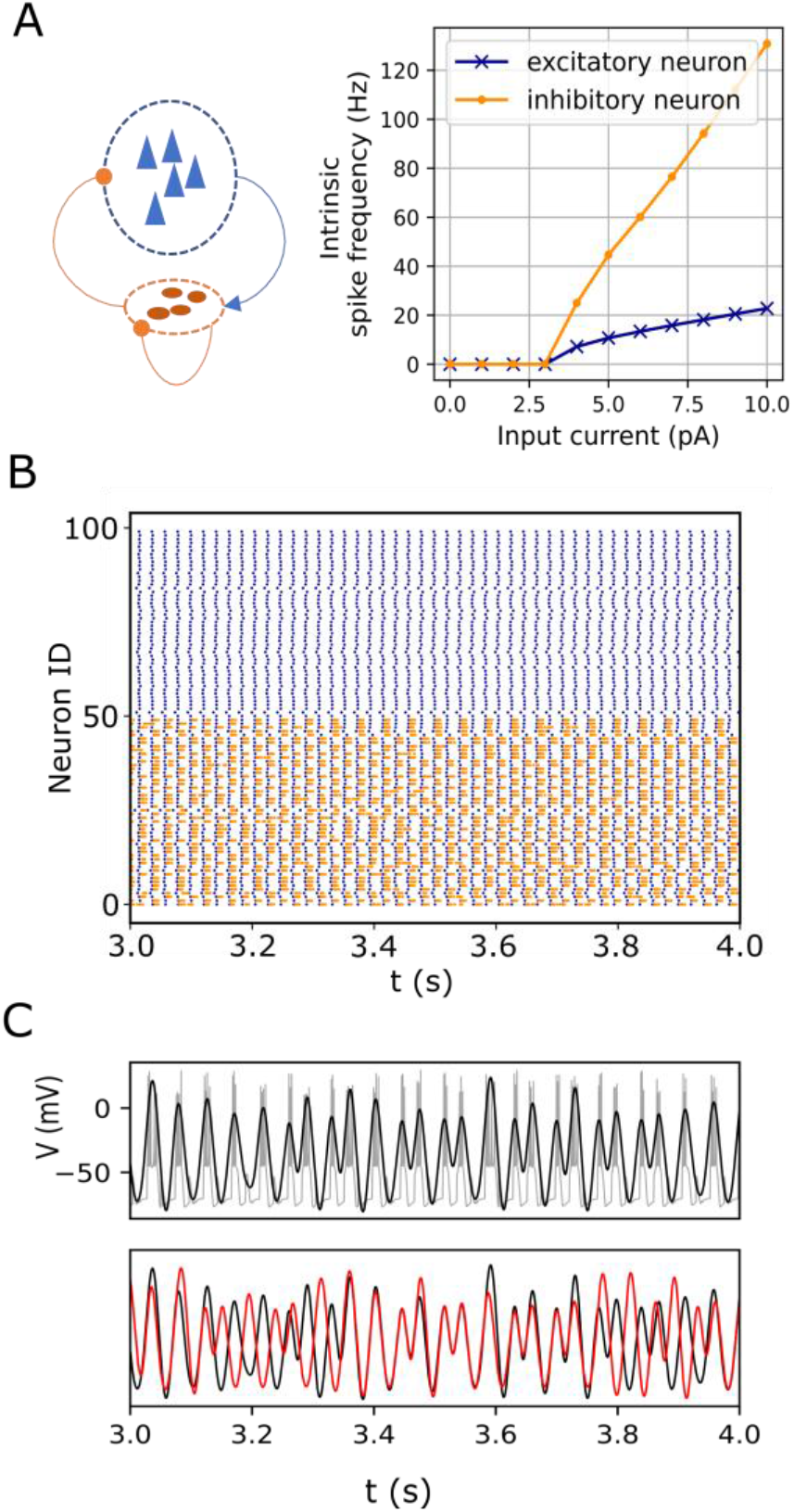
**A.** *Left*: Network configuration consisting of pyramidal neurons (blue triangles) and fast-spiking interneurons (dark orange ovals). Connections indicated by the arrowhead and circles denote excitatory and inhibitory pulse-coupling, respectively. *Right*: Spiking frequencies of an isolated excitatory (pyramidal) neuron and an inhibitory (fast-spiking) neuron for various input currents. **B.** Raster plot showing spiking activity in pyramidal (blue) and fast-spiking neurons (orange) for one second. **C.** (top) Bursting activity (gray) in one representative fast-spiking neuron. The filtered signal, superimposed in black, captures the burst oscillations. (bottom) Burst oscillations of two representative neurons (black and red) showing transitions between in- and anti-phase transiently locked modes.

### 2.1. Assessing the stability of phase relationships

To assess the nature of population-wide phase relationships among the interneurons (our focus of interest), we calculated the phase differences between neuron pairs over time. This resulted in an ensemble of phase difference measurements between all possible distinct pairs of neurons ((50*49)/2=1225 pairs) at each time point over the entire duration (10s). Since time is discrete with a resolution of 0.1ms, this ensemble consisted of N=122,500,000 phase angle difference measurements in total. Henceforth, these angles will be referred to as *ϕ_j_*, *j* = 1,2,…,*N*. We found that the ensemble of angles *ϕ_j_* was organized into clusters. To characterize this, we calculated the Kuramoto-Daido order parameters *Z_n_* [10,17,18]:

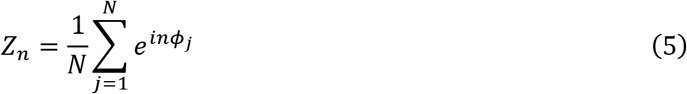

for *n*, a positive integer. These order parameters characterize clustering patterns in the ensemble as follows. If each ensemble member is plotted as a point on the unit circle in the complex plane at its angle, *ϕ_j_*, then *Z*_1_ is the centroid of the ensemble. Of interest is the magnitude |*Z*_1_|, which ranges from zero to one. If all points occur at the same angle, then |*Z*_1_| = 1. In contrast, if the points are scattered uniformly about the unit circle, then |*Z*_1_| = 0. Additionally, if the ensemble of points consists of uniformly spaced clusters, then |*Z*_1_| = 0. Such arrangements can be identified using the order parameters *Z_n_* with *n* > 1. For example, if the phases form three ideal clusters (i.e., equally populated clusters 120 degrees apart), then |*Z*_1_| = |*Z*_2_| = 0, but |*Z*_3_| = 1. In fact, for ideal states consisting of *c* uniformly spaced and equally populated clusters, |*Z_jc_*| = 1 for any integer *j* ≥ 1 and is zero otherwise. To correct for this redundancy, we defined a real-valued quantity *G_n_*

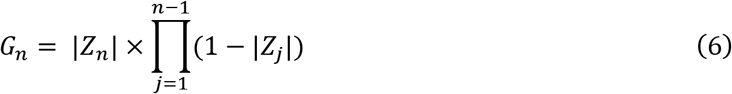

as in Equation 6, such that *G_n_* is close to 1 only when the ensemble approximates an ideal *n*-cluster state.

### 2.2. Characterizing exogenous transitions

To assess the nature of transitions between clustered states in different metastable networks, the following steps were carried out: The networks were simulated twice with identical initializations. The first simulation was used to establish the baseline instantaneous clustering of neurons at any given time (i.e., clustering of neurons based on their instantaneous phases). In the

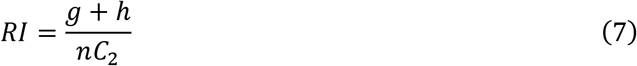

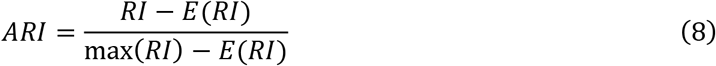

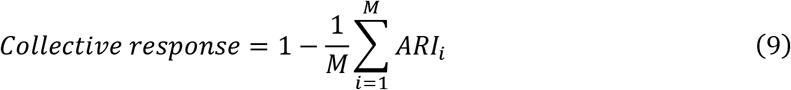

second simulation, a fraction of randomly selected interneurons was simultaneously perturbed by single spikes at a certain timepoint. This caused some of the neurons to change their cluster assignments relative to the baseline. Phase cluster assignments of neurons were obtained using Gaussian mixture models [19]. To quantify the effect of the perturbation, we measured the similarity of the two clustering arrangements of neurons between the baseline and perturbed conditions over time using Equations 7 – 9. Here, *RI* is the “rand index,” which measures the similarity of two sets of instantaneous phase clusters, and *g* and *h* are the numbers of pairs of neurons with the same and different cluster assignments across two groupings (i.e., the number of pairwise agreements between two clustering arrangements). *nC*_2_ denotes the number of possible combinations of neuron pairs. Thus, the *RI* for two identical groupings of a set of neurons is one. The adjusted rand index (ARI) defined in Equation – 8 [20–22] was used to account for chance, and it ensures that the similarity of two random groupings of a set of neurons is zero. This is achieved by estimating an expected RI (*E* (*RI*) in Equation – 8) using random permutations of cluster assignments.

We defined the population’s collective response as in Equation – 9 to characterize the extent to which the network (as a whole) responds to exogeneous perturbations of a fraction of its elements. The ARIs were measured in 50 simulation trials (*M* = 50 in Equation – 9), each with a random subset of perturbed interneurons and a random time (>0.5s) at which the perturbation was delivered. The observed average change in ARI following the perturbation denoted the effect of the perturbation. For instance, if there is no change to the network state following the perturbation, then the collective response is zero. On the other hand, if a perturbation results in new phase clusters that are equivalent to completely random cluster assignments for all the neurons compared to the baseline condition, then the collective response is one. Finally, the magnitude of collective response was examined in metastable networks for various numbers of perturbed neurons. It is worth mentioning here that while each presynaptic spike influences the activity of a postsynaptic neuron (see Equation 4), the collective response aims to characterize activity that is spatiotemporally coarse-grained (i.e., transitions that occur among a group of neurons on a relatively larger timescale than that of the individual spikes).

## 3. Results

The simulated network model incorporates a significant difference in the spiking time scales of the pyramidal and fast-spiking interneurons. The slower excitatory drive from the pyramidal neurons modulates the fast-spiking dynamics of the interneurons, resulting in bursting oscillations in the latter (**Fig. 1B&C**). The inhibitory interactions between these bursting neurons then leads to itinerant metastable behavior. That is, ongoing endogenous transitions occur between different phase-locked (attractor) states. An example is illustrated in **Fig. 1C**, which shows the bursting oscillations of a representative pair of neurons versus time. At first, the neurons are locked in phase, but then they transition to an out-of-phase state. After a short time, they return to the in-phase state, and then switch back to the out-of-phase state. Similar persistent behavior was described in a previous report [10] (also see Supplementary **Fig. S1**).

### 3.1. Frequency dependence of phase cluster stability

The frequency of pyramidal neuron spiking is determined by the value of *I^ext^*. To examine the effect of this driving frequency on the characteristics of metastability in the fast-spiking interneuron network, we systematically varied *I^ext^* and measured the distribution of phase differences (*ϕ_j_*) between pairs of interneurons as described in Methods **Section 2.1**. **Fig. 2** shows a plot of *G_n_* (*n*=1, 2,…,7), which quantifies the degree to which the interneuron phase differences group into *n* clusters, versus the pyramidal neuron spiking frequency (*f*). Above the plot are polar histograms of the phase differences (*ϕ_j_*) for selected *f*’s. At low driving frequency, all the inhibitory neurons lock in-phase, and thus their phase differences are close to zero. Correspondingly, *G*_1_ is high and *G*_2_ through *G*_7_ are low. Interestingly, as the driving frequency is increased beyond ~25*Hz*, clusters appear in the phase difference distribution. We find that a well-defined state of two clusters emerges near *f* =48Hz, and a three-cluster state emerges near *f* =77Hz. As the driving frequency is further increased, the pattern reverses and another well-defined state of two clusters appears near *f* =117Hz. Beyond *f* =140Hz, all *G_n_* are small and the phase difference distributions become more uniform. It is interesting to note that the emergence of these clustered states occurs for driving frequencies within the gamma range. Pyramidal neurons are indeed known to fire at gamma frequencies, although such gamma cycles have been observed to be modulated by slower theta cycles [23,24]. These results show that the metastable attractors emerge in an interneuron population when their spiking dynamics are appropriately driven by slower excitatory frequencies.

**Figure 2.**
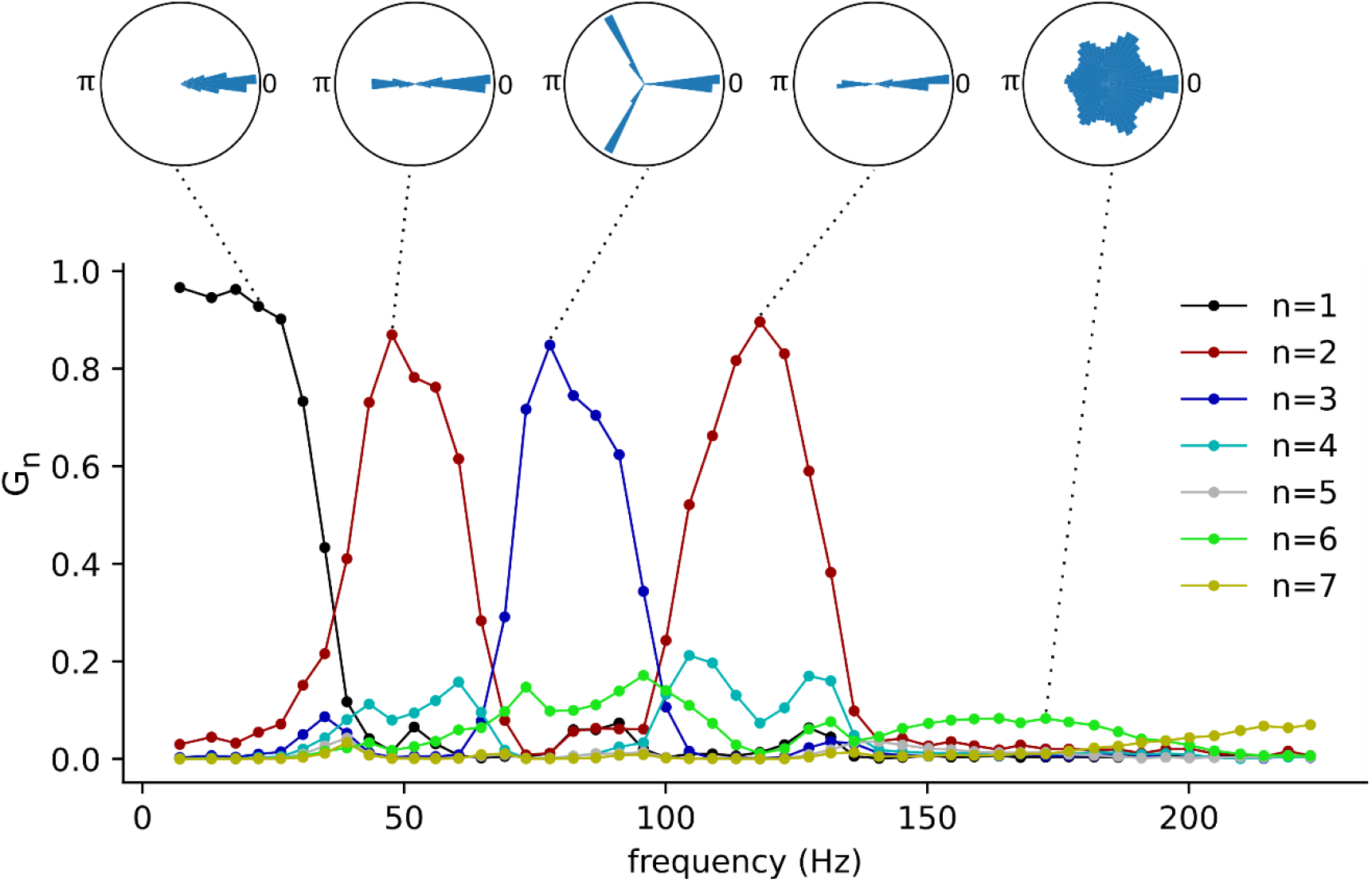
The network-wide stability and the number of phase clusters in the fast-spiking interneurons as functions of pyramidal neuron frequency. Polar histograms show phase differences *ϕ_j_* for all pairs of neurons in representative networks. Pyramidal neuron frequency was computed by taking the inverse of the average population interspike intervals.

The phase-clustering phenomena described above were consistently observed for a range of connection probabilities (**Fig. 3**). We analyzed networks instantiated on a 50×50 grid of inter- and intra-group connection probabilities in the range [0, 1]. The *G*_2_ and *G*_3_ versus frequency curves were computed as described before for each of the 2500 networks. **Fig. 3A** shows the maximum of *G*_2_ (left) and *G*_3_ (right) across frequencies for each network. These measures were sensitive to the inter-group connection probabilities and relatively less sensitive to the intra-group connection probabilities. Plotting these measures against the frequency of excitatory neurons (**Fig. 3B**) revealed that the stable phase clusters emerged approximately within the range [25*Hz*, 200*Hz*], Our results suggest that the emergent attractors’ stability in interneurons depend on their connectivity with excitatory neurons and the frequency of excitatory oscillations. There are robust gradients towards maximally stable interneuron phase cluster arrangements along the dimensions of pyramidal frequency and E-I connection probability (**Figs. 2&3**).

**Figure 3.**
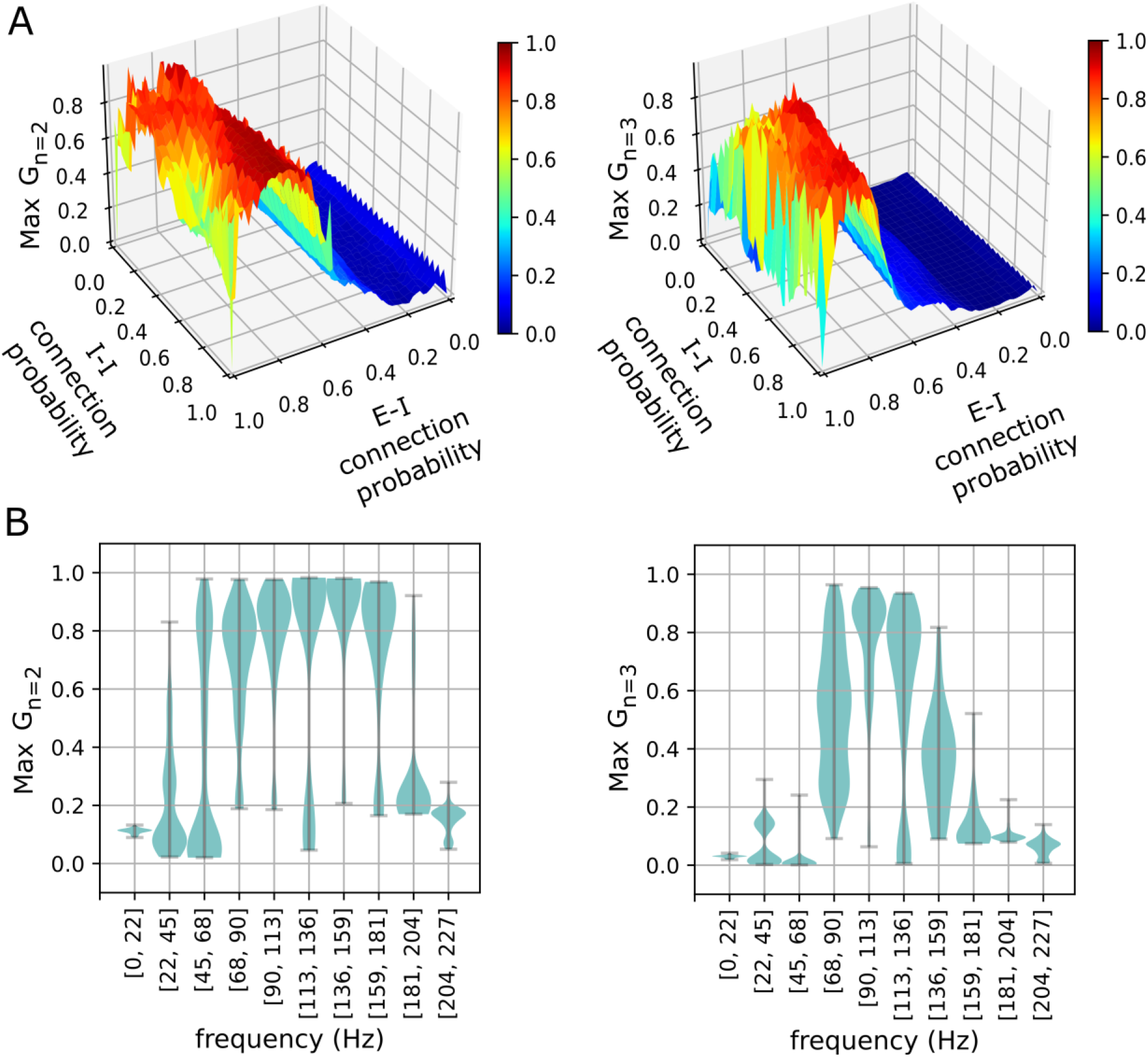
The stability of phase clusters against network connectivity and pyramidal neuron frequency. **A.** The most stable two-cluster (left) and three-cluster (right) states realized in networks for various excitatory-inhibitory (E-I) and inhibitory-inhibitory (I-I) connection probabilities. **B.** Distributions of maximum phase cluster stability are plotted against their respective frequency bins of pyramidal neurons for two-cluster (left) and three-cluster (right) states. Width indicates the density of points for a given maximum stability.

### 3.2. Collective response characteristics of metastable networks

The multi-cluster states described above exhibit spontaneous and endogenous itinerant dynamics (**Fig. 1C** and Supplementary **Fig. S1**). Motivated by the fact that an organism adapts its internal state to the environment via external stimuli, we sought to measure how a metastable state would be affected by an external perturbation. As described in Methods **Section 2.2**, the network was simulated twice with identical initializations. The first simulation established the baseline dynamics as shown in **Fig. 4A** (left, all rows). The panels on the left show the instantaneous phases of individual neurons in the interneuron population at three instants of time. The neurons are ordered along the horizontal axis by increasing phase. To mask the endogenous itinerant behavior (see Supplementary **Fig. S1)**, the neurons were reordered for each left panel in **Fig. 4A**. In the second simulation, an external perturbation was applied. Five randomly selected interneurons each received a single spike at time *t_p_*. The results are shown in the right panels of **Fig. 4A**. The time of perturbation occurred between the top and middle panels. Note that the neuron orderings from the left columns are preserved in the right columns. Thus, any differences in the cluster assignment of neurons are due exclusively to the perturbation. Additionally, the transitions are collective in the sense that they occur in many non-perturbed neurons immediately following the perturbation (**Fig. 4A** - right). This highlights the synergy of interacting elements in metastable networks. Collective transitions may be characterized by treating network states as multiple instantaneous phase clusters of neurons at any given time and subsequently comparing the cluster assignments of neurons between unperturbed and perturbed conditions using the ARI (**Fig. 4B**). For example, when *t* < *t_p_*, the two simulation conditions (left and right panels of **Fig. 4A**) show identical cluster arrangements of neurons resulting in ARI values of 1. However, immediately following perturbation (*t* > *t_p_*), the network states begin to diverge between the two simulation conditions resulting in ARI values less than 1 (**Fig. 4B**).

**Figure 4.**
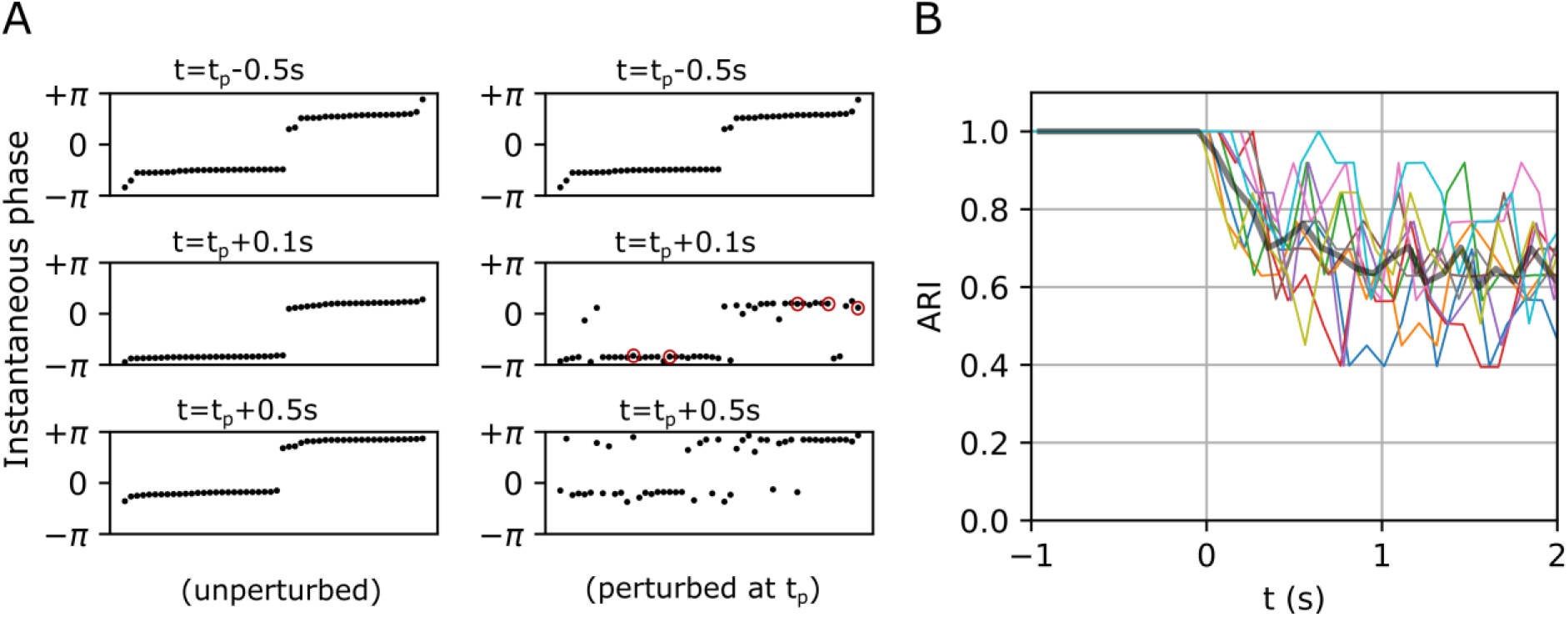
Exogenous transitions in the fast-spiking interneuron population. **A.** Left and right panels show the temporal evolution of a network in unperturbed and perturbed conditions respectively. Neurons (black dots) in the left panels are sorted by their instantaneous phases at each *t*, and their order is preserved in the right panels to show the divergence of states caused by perturbation. *t_p_* denotes the time of perturbation, and red circles in the right middle panel denote perturbed neurons. **B.** Ten representative ARI curves (color) computed by comparing cluster assignments of neurons between unperturbed and perturbed conditions. The black curve represents the average. Here, t=0 corresponds to the perturbation times *t_p_*.

**Fig. 5** shows the magnitude of collective responses (Equation – 9) in three representative metastable networks that showed high magnitudes of stability in **Fig. 2**. The two-cluster states that emerged near *f* =48Hz in Fig. 2 (see Section 3.1) showed the strongest response, reaching magnitudes of 0.4 within 1s when sufficient number of neurons were perturbed (**Fig. 5A**). On the other hand, the three-cluster states, which emerged near *f* =77Hz, elicited slower responses to external perturbations (**Fig. 5B**), and the two-cluster states that emerged near *f* =117Hz elicited the slowest response (**Fig. 5C**). Note that *f* directly affects the inter-burst intervals in interneurons and consequently shapes their multi-periodic trajectories in the phase space. Taken together, these results suggest that the metastability realized in interneuron populations may arise from mechanisms [13,25,26] that evolve depending on (a) their connectivity with excitatory neurons and (b) the frequencies of excitatory oscillations within a cortical region.

**Figure 5.**
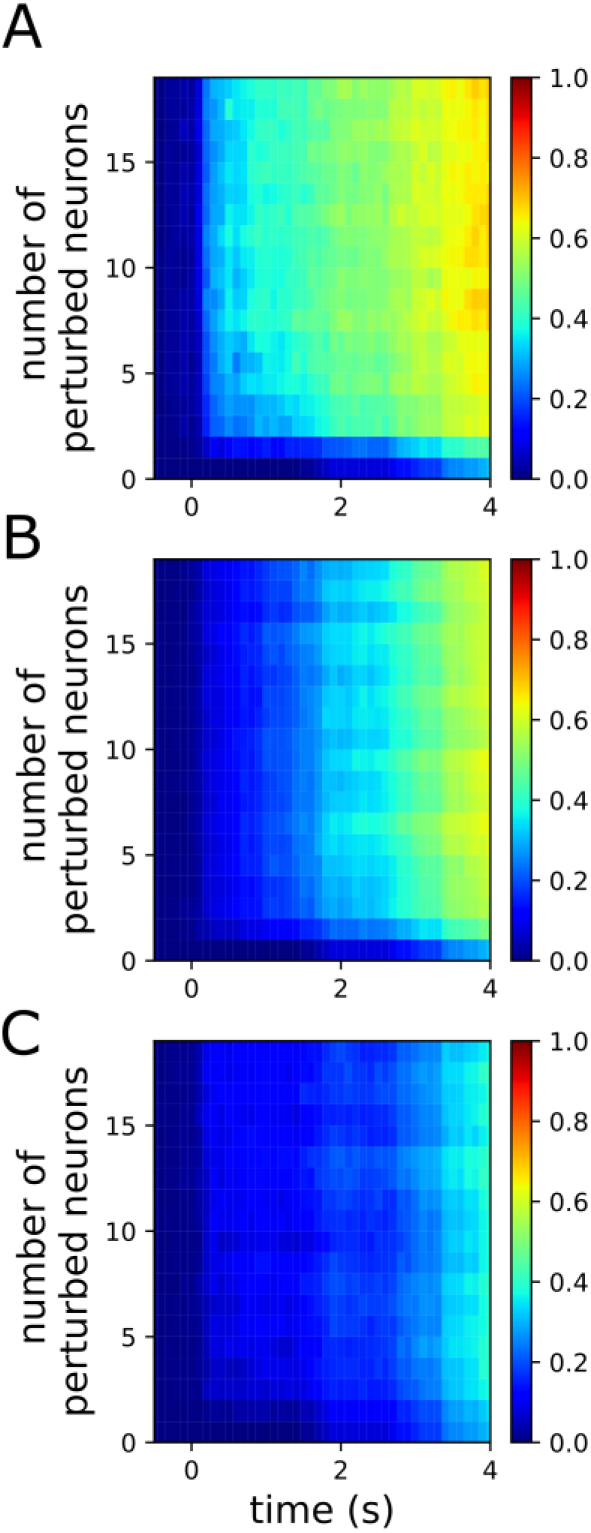
Magnitude of collective response (equation – 9) characterized by the exogenous transitions in three representative networks for various numbers of perturbed neurons. The pyramidal driving frequencies used are 48Hz (A), 77Hz (B), and 117Hz (C); these correspond to two, three, and two cluster states in the interneuron population, respectively (see Fig. 2).

## 4. Discussion

The modern examination of neural connectivity has enjoyed considerable attention through several early key theoretical studies [27–30]. Along these lines, understanding the emergent dynamical complexities in a network in terms of its constituent mechanisms remains an important goal. Here, we studied the emergent phenomenon of metastability in spiking neural network models that represent cortical local circuits. Metastability is characterized by the coexistence of integrated (synchronized states) and segregated (transitions between synchronized states) behaviors in a system of interacting elements [11]. We specifically studied how the neuron-level spiking frequencies are related to the emergent metastability, and how these relationships are governed by the underlying excitatory and inhibitory neuronal connectivity. Moreover, the spiking neural network models examined in this paper provide compact mesoscopic scale approximations of regional collective dynamics that capture a crucial biological complexity at the level of individual neurons. A limitation of the current work is that it only considered two types of neuron populations, whereas intraregional neural circuits consist of populations with diverse frequency profiles [31]. Nevertheless, current work demonstrates that the emergent metastability in an interneuron population depends on their connectivity with excitatory neurons and the frequency of excitatory drive. Their relationships are revealed in the *G_n_* curves that show robust gradients along the dimensions excitatory-inhibitory connection probability (**Fig. 3A**) and the frequency of the excitatory drive (**Figs. 2 & 3B**). Thus, a numerical optimization using *G_n_* as an objective function can, in principle, estimate optimal intraregional connectivity configurations among groups of neurons with heterogenous frequency profiles.

Additionally, the models and the analyses methods presented in this paper are easily scalable to study interregional interactions in the brain. Empirically estimated anatomical connectivity between brain regions provides useful information for modeling interregional metastability. Note that while the current work suggests a link between two temporal scales (i.e., timescales of neuronal spiking and attractor transitions) of network dynamics, modeling interregional interactions using their anatomical connectivity will further provide a venue to study metastable brain dynamics at a macroscopic spatial scale. In the next section, we present a design for whole-brain modeling and discuss biological observations of atypical brain physiology that can be mechanistically interpreted in such models.

### 4.1. A blueprint for whole-brain modeling at the mesoscopic level

Sustained background activity in the cortex arises from the mutual excitation of excitatory neurons [32] such as the pyramidal neurons that project their axons to longer distances than the interneurons. Such long-distance projections are captured in diffusion-weighted magnetic resonance imaging (dMRI) as white matter tracts. Estimation of anatomical connectivity from dMRI relies on the tractography-derived streamline counts between regions [33]. However, the direction of axonal projections is not identifiable from the dMRI tractography. In other words, if the tractography estimates that there are *n* streamlines between regions A and B, the fraction of *n* that corresponds to the axonal projections from region A to B (and vice versa) is unidentifiable based on the tractography alone. The net excitatory input to a region is a key determinant of its firing frequency. Since the regional metastable dynamics may depend upon the frequency of excitatory oscillations (See **Figs. 2&5**), we consider that resolving the directionality of fiber tracts (i.e., incoming vs. outgoing for a region) will be important to accurately model the whole-brain metastable dynamics.

Previous whole-brain modeling studies considered symmetric couplings between pairs of brain regions and estimated a global coupling constant to supplement the connectivity information derived from white matter tractography [34,35]. The unknown degrees of asymmetries in structural connections could be estimated for the whole brain, in principle, by numerical optimization. The network-wide stability of attractors (Equation – 6) realized across the frequency spectrum in whole-brain models could serve as a heuristic metric to search for the directionality, or more generally, the asymmetries in the bidirectionality, of macroscopic anatomical connections. Note that the number of unknown parameters is directly related to the spatial resolution of structural connectivity specification. It is uncertain if a model description at a high spatial resolution is solvable in practice due to the computational demands. However, models can be described at various spatial resolutions of a group-averaged whole-brain connectome, and the convergence patterns of optimization solutions for each spatial resolution can be examined. These convergence patterns should be compared against those of the null models with randomly shuffled anatomical connections. The robustness of the solution convergence relative to that of the respective null model can inform the preferred spatial resolution for the whole-brain models. This approach considers that the brain-wide features of the human connectome evolved to be highly specific, and they provide strong constraints to model network-wide transient attractor dynamics. The identified directionalities can also be validated against findings from the morphological tracing studies that report the directions of axonal projections between cortical regions using anterograde or retrograde tracing techniques [36].

The whole-brain models of anatomically constrained mesoscopic scale metastable dynamics are useful mathematical tools that will enable a multiscale understanding of cortical dynamics and illuminate neural network mechanisms of pathophysiology in neurological disorders. As an example, we briefly discuss how a whole-brain model of metastable dynamics could be useful in elucidating the pathophysiology of Parkinson’s disease (PD). The structural degeneration of cortical and subcortical connectivity is a key factor underlying the motor and cognitive deficits in PD [37–39]. However, the mechanisms by which the structural degeneration alters the brain-wide dynamics in PD is not well understood. A whole-brain model of metastable dynamics constrained by the PD-affected anatomical connectivity can potentially delineate such mechanisms. For instance, amplified synchronization in beta frequency (8Hz – 30Hz), which has been reported in the cortical and basal ganglia circuits of PD subjects, is hypothesized to contribute to PD symptoms [37,40,41]. In particular, longer episodes of beta oscillations in the subthalamic nucleus (STN) [42] and its higher synchronization with the cortical supplementary motor area (SMA) were associated with the freezing of gait in PD [43]. It may be hypothesized that the freezing of gait in PD is contributed by a reduction in synergistic transitions between different attractor states realized in these areas. The increased beta synchronization between STN and SMA observed in PD, and the hypothesized reduction in attractor transitions can be validated in whole-brain models by examining their stability (Equation – 6) and collective response characteristics (Equation – 9) respectively. Furthermore, by selectively targeting frequency bands for manipulation, one can investigate causal relationships between specific frequencies of oscillations and the dynamics of transitions between attractor states realized across multiple timescales. Such whole-brain models could also enable identification of network mechanisms to restore optimal brain dynamics in PD and other neurological disorders.

## 5. Conclusion

The numerical simulations in this study show that the inhibitory interneuron oscillations that are modulated by slower excitatory neurons, under certain conditions, realize metastable attractors. Furthermore, our analyses revealed frequency dependences of the emergent metastability and quantitatively associated the features of spiking neural network dynamics at two temporal scales. This novel cross-timescale association using a compact mathematical description captures a biological complexity of neural circuits at the level of individual neurons and allows, in principle, a scalable integration with empirical observations of neural network features to develop a coherent understanding of nervous system functions.

## Acknowledgments

The authors thank Alev Erisir, Teague Henry, Giselle Petzinger, Michael Jakowec, and Dawn Schiehser for their insights and helpful discussions. The authors appreciate the complementary perspectives of Jeffrey Kopsick, Giorgio Ascoli, and Alexander Komendantov and are thankful for their many valuable discussions.

## Author Contributions

SV, EB and JDVH conceptualized the study. SV and AS contributed to the analyses. EB and HS provided critical inputs at various stages of this study. SV and EB contributed to manuscript writing. EB and JDVH contributed to supervision and JDVH contributed to funding acquisition.

## Data Availability

The data that support the findings were generated from numerical simulations by the authors, and all simulation and analysis scripts are publicly available at *https://github.com/sivaven/Transient-Synchronization*.

