## Supplementary Fig. S1 for "A spiking neural network model of cortical intraregional metastability"

### Supplementary material

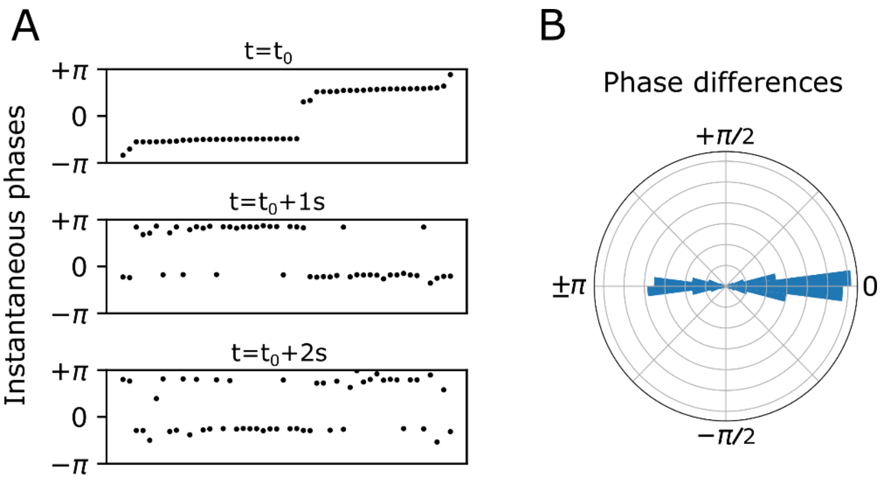


**Supplementary Figure S1. A.** Endogenous itinerant states characterized by phase clusters at three time points over a 2s duration. Neurons (black dots) are ordered by their instantaneous phases in the top panel, and this ordering is preserved in the bottom two panels. Two phase clusters persist over time, and individual neurons spontaneously switch from one cluster to another. **B.** Persistent phase relationships between neurons are illustrated in a polar histogram by plotting the phase differences between all distinct pairs of neurons over the 2s duration.
